# Genome-wide analyses supported by RNA-Seq reveal non-canonical splice sites in plant genomes

**DOI:** 10.1101/428318

**Authors:** Boas Pucker, Samuel F. Brockington

**Affiliations:** Evolution and Diversity, Department of Plant Sciences, University of Cambridge, Cambridge, United Kingdom; Genetics and Genomics of Plants, CeBiTec & Faculty of Biology, Bielefeld University, Bielefeld, Germany

**Keywords:** gene structure, splicing, annotation, comparative genomics, transcriptomics, gene expression, natural diversity, evolution

## Abstract

Most eukaryotic genes comprise exons and introns thus requiring the precise removal of introns from pre-mRNAs to enable protein biosynthesis. U2 and U12 spliceosomes catalyze this step by recognizing motifs on the transcript in order to remove the introns. A process which is dependent on precise definition of exon-intron borders by splice sites, which are consequently highly conserved across species. Only very few combinations of terminal dinucleotides are frequently observed at intron ends, dominated by the canonical GT-AG splice sites on the DNA level.

Here we investigate the occurrence of diverse combinations of dinucleotides at predicted splice sites. Analyzing 121 plant genome sequences based on their annotation revealed strong splice site conservation across species, annotation errors, and true biological divergence from canonical splice sites. The frequency of non-canonical splice sites clearly correlates with their divergence from canonical ones indicating either an accumulation of probably neutral mutations, or evolution towards canonical splice sites. Strong conservation across multiple species and non-random accumulation of substitutions in splice sites indicate a functional relevance of non-canonical splice sites. The average composition of splice sites across all investigated species is 98.7% for GT-AG, 1.2% for GC-AG, 0.06% for AT-AC, and 0.09% for minor non-canonical splice sites. RNA-Seq data sets of 35 species were incorporated to validate non-canonical splice site predictions through gaps in sequencing reads alignments and to demonstrate the expression of affected genes. We conclude that *bona fide* non-canonical splice sites are present and appear to be functionally relevant in most plant genomes, if at low abundance.

## Introduction

Introns separate eukaryotic genes into exons [1, 2]. After their likely origin as selfish elements [3], introns subsequently evolved into beneficial components in eukaryotic genomes [4–6]. Historical debates concerning the evolutionary history of introns led to the “introns-first-hypothesis” which proposes that introns were already present in the last common ancestor of all eukaryotes [3, 7]. Although this putative ancestral genome is inferred to be intron-rich, several plant genomes accumulated more introns during their evolution generating the highly fragmented gene structures with average intron numbers between six and seven [8]. Introner elements (IEs) [9], which behave similar to transposable elements, are one possible mechanism for the amplification of introns [10]. Early introns probably originated from self-splicing class II introns [3, 11] and evolved into passive elements, that require removal by eukaryote-specific molecular machineries [11]. No class II introns were identified in the nuclear genomes of sequenced extant eukaryotes [11] except for mitochondrial DNA (mtDNA) insertions [12, 13].

The removal of these introns during pre-mRNA processing is a complex and expensive step, which involves 5 snoRNAs and over 150 proteins building the spliceosome [14]. In fact, a major U2 [15] and a minor U12 spliceosome [16] are removing different intron types from eukaryotic pre-mRNAs [17]. The major U2 spliceosome mostly recognises canonical GT-AG introns, but is additionally reported to remove AT-AC class I introns [18]. Non-canonical AT-AC class II introns are spliced by the minor U2 spliceosome, which is also capable of removing some GT-AG introns [18, 19]. Highly conserved cis-regulatory sequences are required for the correct spliceosome recruitment to designated splice sites [20–22]. Although these sequences pose potential for deleterious mutations [4], some intron positions are conserved between very distant eukaryotic species like *Homo sapiens* and *Arabidopsis thaliana* [23].

Among the most important recognition sequences of spliceosomes are dinucleotides at both ends of spliceosomal introns which show almost no variation from GT at the 5’ end and AG at the 3’ end, respectively [24]. Different types of alternative splicing generate diversity at the transcript level by combining exons in different combinations [25]. This process results in a substantially increased diversity of peptide sequences [2, 26]. Special splicing cases e.g. utilizing a single nucleotide within an intron for recursive splicing [27] or generating circular RNAs [28] are called non-canonical splicing events [25] and build an additional layer of RNA and proteomic diversity. If this process is based on splice sites differing from GT-AG those splice sites are called non-canonical. Non-canonical splice sites were first identified before genome sequences became available on a massive scale (reviewed in [29]). GC-AG and AT-AC are classified as major non-canonical splice site combinations, while all deviations from these sequences are deemed to be minor non-canonical splice sites. More recently, advances in sequencing technologies and the development of novel sequence alignment tools now enable a systematic investigation of non-canonical splicing events [25, 30]. Comprehensive genome sequence assemblies and large RNA-Seq data sets are publicly available. Dedicated split-read aligners like STAR [31, 32] are able to detect non-canonical splice sites during the alignment of RNA-Seq reads to genomic sequences. Numerous differences in annotated non-canonical splice sites even between accessions of the same species [30] as well as the extremely low frequency of all non-canonical splice sites indicate that sequencing, assembly, and annotation are potential major sources of erroneously inferred splice sites [29, 30, 33]. Distinguishing functional splice sites from degraded sequences such as in pseudogenes is also still an unsolved issue. Nonetheless, the combined number of currently inferred minor non-canonical splice site combinations is even higher than the number of the major non-canonical AT-AC splice site combinations [30, 34].

Here, we analysed 121 whole genome sequences from across the entire plant kingdom to harness the power of a very large sample size and genomic variation accumulated over extensive periods of evolutionary time, to better understand splice site combinations. Although, only a small number of splice sites are considered as non-canonical, the potential number in 121 species is large. Furthermore, conservation of sequences between these species over a long evolutionary time scale may also serve as a strong indication for their functional relevance. We incorporated RNA-Seq data to differentiate between artifacts and *bona fide* cases of active non-canonical splice sites. Active splice sites are revealed by an RNA-Seq read alignment allowing quantification of splice site activity. We then identified homologous non-canonical splice sites across species and subjected the genes containing these splice sites to phylogenetic analyses. Conservation over a long evolutionary time, expression of the effected gene, and RNA-Seq reads spanning the predicted intron served as evidence to identify *bona fide* functional non-canonical splice site combinations.

## Materials & Methods

### Collection of data sets and quality control

Genome sequences (FASTA) and the corresponding annotation (GFF3) of 121 plant species (Additional file 1) were retrieved from the NCBI. Since all annotations were generated by GNOMON [35], these data sets should have an equal quality and thus allow comparisons between them. BUSCO v3 [36] was deployed to assess the completeness and duplication level of all sets of representative peptide sequences using the reference data set ‘embryophyta odb9’.

### Classification of annotated splice sites

Genome sequences and their annotation were processed by a Python script to identify the representative transcript per gene defined as the transcript that encodes the longest polypeptide sequence [30, 37]. Like all custom Python scripts relevant for this work, it is available with additional instructions at https://github.com/bpucker/ncss2018. Genes with putative annotation errors or inconsistencies were filtered out as done before in similar analyses [38]. Focusing on the longest peptide is essential to avoid biases caused by different numbers of annotated isoforms in different species. Splice sites within the coding sequence of the longest transcripts were analyzed by extracting dinucleotides at the borders of all introns. Untranslated regions (UTRs) were avoided due to their more challenging and thus less reliable prediction [30, 39]. Splice sites and other sequences will be described based on their encoding DNA sequence (e.g. GT instead of GU for the conserved dinucleotide at the donor splice site). Based on terminal dinucleotides in introns, splice site combinations were classified as canonical (GT-AG) or non-canonical if they diverged from the canonical motif. A more detailed classification into major non-canonical splice site combinations (GC-AG, AT-AC) and all remaining minor non-canonical splice site combinations was applied. All following analyses were focused on introns and intron-like sequences equal or greater than 20 bp.

### Investigation of splice site diversity

A Python script was applied to summarize all annotated combinations of splice sites that were detected in a representative transcript. The specific profile comprising frequency and diversity of splice site combinations in individual species was analyzed. Splice site combinations containing ambiguity characters were masked from this analysis as they are most likely caused by sequencing or annotation errors. Spearman correlation coefficients were computed pairwise between the splice site profiles of two species to measure their similarity. Flanking sequences of CA-GG and GC-AG splice sites in rice were investigated, because CA-GG splice sites seemed to be the result of an erroneous alignment. The conservation of flanking sequences was illustrated based on sequence web logos constructed at https://weblogo.berkeley.edu/logo.cgi.

### Analysis of splice site conservation

Selected protein encoding transcript sequences with non-canonical splice sites were subjected to a search via BLASTn v2.2.28+ [40] to identify homologues in other species to investigate the conservation of splice sites across plant species. As proof of concept, one previously validated non-canonical splice site containing gene [30], At1g79350 (rna15125), was investigated in more depth. Homologous transcripts were compared based on their annotation to investigate the conservation of non-canonical splice sites across species. Exon-intron structures of selected transcripts were plotted by a Python script using matplotlib [41] to facilitate manual inspection.

### Validation of annotated splice sites

Publicly available RNA-Seq data sets of different species (Additional file 2) were retrieved from the Sequence Read Archive [42]. Whenever possible, samples from different tissues and conditions were included. The selection was restricted to paired-end data sets to provide a high accuracy during the read mapping. Only species with multiple available data sets were considered for this analysis. All reads were mapped via STAR v2.5.1b [31] in 2-pass mode to the corresponding genome sequence using previously described cutoff values [43]. A Python script utilizing BEDTools v2.25.0 [44] was deployed to convert the resulting BAM files into customized coverage files. Next, the read coverage depth at all exon-intron borders was calculated based on the terminal nucleotides of an intron and the flanking exons. Splice sites were considered as supported by RNA-Seq if the read coverage depth dropped by at least 20% when moving from an exon into an intron (Additional file 3).

### Phylogenetic tree construction

RbcL (large RuBisCO subunit) sequences of almost all investigated species were retrieved from the NCBI for the construction of a phylogenetic tree. MAFFT v.7 [45] was deployed to generate an alignment which was trimmed to a minimal occupancy of 60% in each alignment column and finally subjected to FastTree v.2.1.10 [46] for tree construction. Species without an available RbcL sequence were integrated manually by constructing subtrees based on scientific names via phyloT (https://phylot.biobyte.de/). Due to these manual adjustments, the branch lengths in the resulting tree are not accurate and only the topology (Additional file 4) was considered for further analyses.

### Intron length analyses

Stress-related gene IDs of *A. thaliana* were retrieved from the literature [47] and corresponding genes in the NCBI annotations were identified through reciprocal best BLAST hits as previously described [48]. Lengths of introns in these stress genes were compared against an equal number of randomly selected intron lengths from all remaining genes using the Wilcoxon test as implemented in the Python module scipy. Average values of the stress gene intron lengths as well as the randomly selected intron lengths were compared. This random selection and the following comparison were repeated 100 times to correct for random effects.

Minor non-canonical splice site combinations without ambiguous bases in introns longer than 5kb were counted and compared against their frequency in shorter introns. After ranking all splice site combinations by this ratio, the frequency of the four bases A, C, G, and T was analyzed in correlation to their position in this list.

### Comparison of non-canonical splice sites to overall sequence variation

A previously generated variant data set [48] was used to identify the general pattern of mutation and variant fixation between the two *A. thaliana* accessions Columbia-0 and Niederzenz-1. All homozygous SNPs in a given VCF file were considered for the calculation of nucleotide substitution rates. Corresponding substitution rates were calculated for all minor non-canonical splice sites by assuming they originated from the closest sequence among GT-AG, GC-AG, and AT-AC. General substitution rates in a species were compared against the observed substitution in minor non-canonical splice sites via Chi^2^ test.

## Results

### Genomic properties of plants and diversity of non-canonical splice sites

Comparison of all genomic data sets revealed an average GC content of 36.3%, an average percentage of 7.8% of protein encoding sequence, and on average 95.7% of complete BUSCO genes (Additional file 5). Averaged across all 121 genomes, a genome contains an average of 27,232 genes with 4.5 introns per gene. The number of introns per gene was only slightly reduced to 4.15 when only introns enclosed by coding exons were considered for this analysis.

Our investigation of these 121 plant genome sequences revealed a huge variety of different non-canonical splice site combinations (Additional file 6, Additional file 7). Nevertheless, most of all annotated introns display the canonical GT-AG dinucleotides at their borders. Despite the presence of a huge amount of non-canonical splice sites in almost all plant genomes, the present types and the frequencies of different types show a huge variation between species (Additional file 8). A phylogenetic signal in this data set is weak if it is present at all. The total number of splice site combinations ranged between 1,505 (*Bathycoccus prasinos*) and 372,164 (*Brasssica napus*). Algae displayed a very low number of minor non-canonical splice site combinations, but other plant genome annotations within land plants also did not contain any minor non-canonical splice site combinations without ambiguity characters e.g. *Medicago truncatula*. *Camelina sativa* displayed the highest number of minor non-canonical splice site combinations (2,902). There is a strong correlation between the number of non-canonical splice site combinations and the total number of splice sites (Spearman correlation coefficient=0.53, p-value=5.5*10^-10^). However, there is almost no correlation between the number of splice sites and the genome size (Additional file 9).

### Non-canonical splice sites are likely to be similar to canonical splice sites

There is a negative correlation between the frequency of non-canonical splice site combinations and their divergence from canonical sequences (r= -0.4297 p-value=7e-13; Fig.1;Additional file 7). Splice sites with one difference to a canonical splice site are more frequent than more diverged splice sites. A similar trend can be observed around the major non-canonical splice sites AT-AC (Fig.2) and the canonical GT-AG. Comparison of the overall nucleotide substitution rate in the plant genome and the divergence of minor non-canonical splice sites from canonical or major non-canonical splice sites revealed significant differences (p-value=0, Chi^2^ test). For example, the substitutions of A by C and A by G were observed with a similar frequency at splice sites, while the substitution of A by G is almost three times as likely as the A by C substitution between the *A. thaliana* accessions Col-0 and Nd-1.

**Fig. 1:**
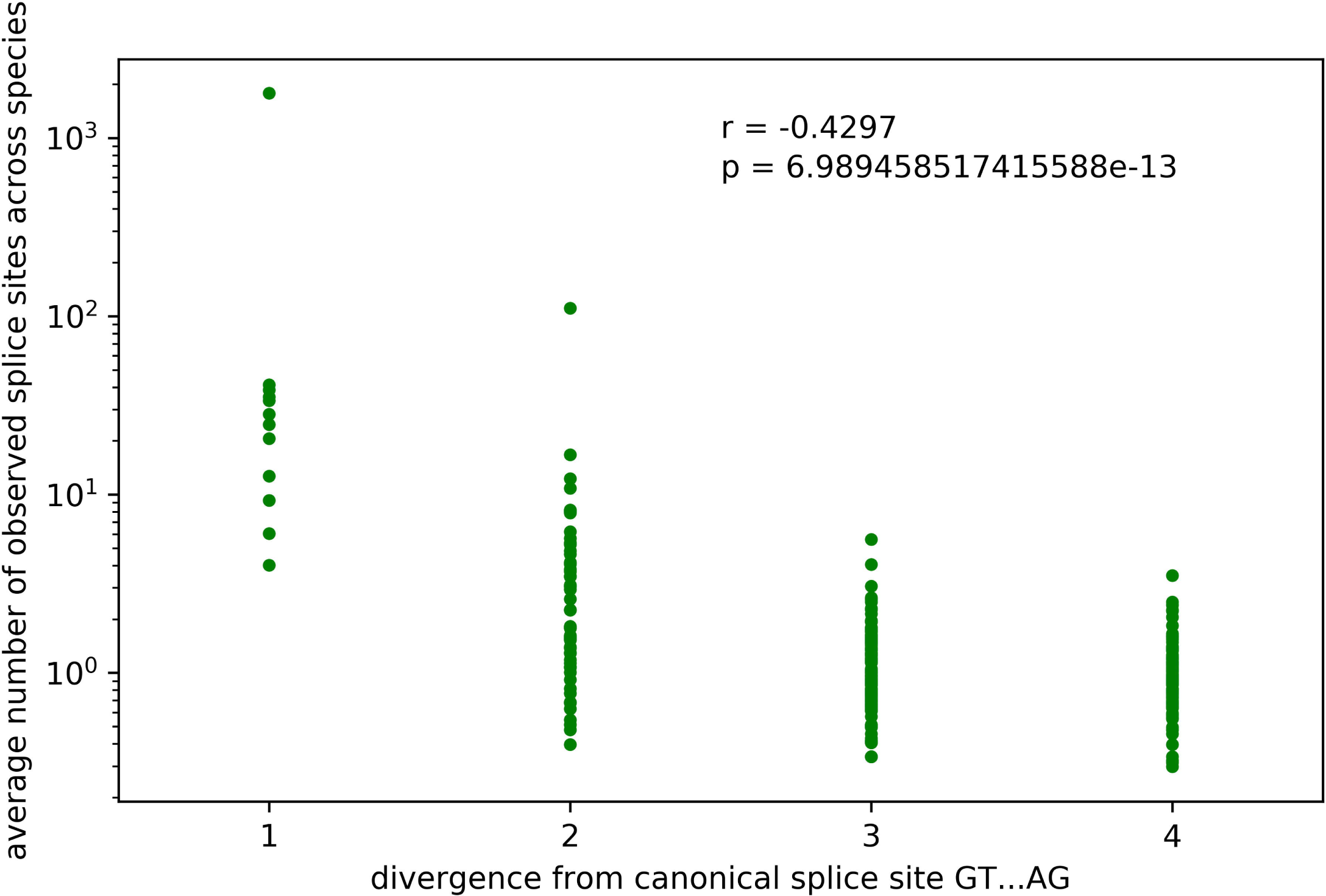
Correlation between splice site sequence divergence and frequency. Spearman correlation coefficient between the splice site combination divergence from the canonical GT-AG and their frequency is r=-0.4297 (p-value = 7*10^-13^).

**Fig. 2:**
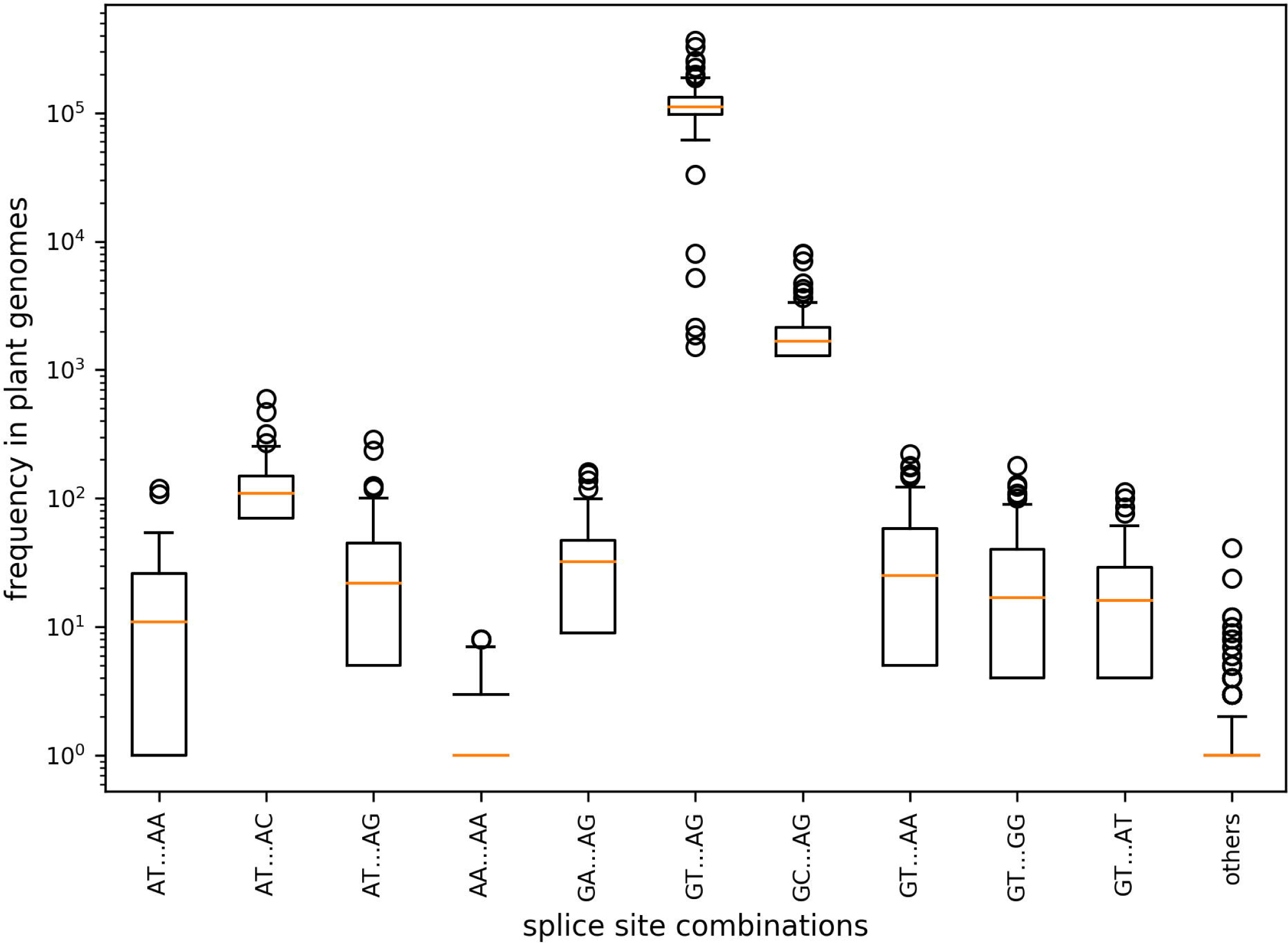
Splice site combination frequency. The frequencies of selected splice site combinations across 121 plant species are displayed. Splice site combinations with high similarity to the canonical GT-AG or the major non-canonical GC-AG/AT-AC are more frequent than other splice site combinations.

The genome-wide distribution of genes with non-canonical splice sites did not reveal striking patterns. When looking at the chromosome-level genome sequences of *A. thaliana, B. vulgaris*, and *V. vinifera* (Additional file 10, Additional file 11, Additional file 12), there were slightly less genes with non-canonical splice sites close to the centromere. However, the total number of genes was reduced in these regions as well, so likely correlated with genic content.

One interesting species-specific property was the high frequency of non-canonical CA-GG splice site combinations in *Oryza sativa* which is accompanied by a low frequency of the major non-canonical GC-AG splice sites. In total, 233 CA-GG splice site combinations were identified. However, the transcript sequences can be aligned in a different way to support GC-AG sites close to and even overlapping with the annotated CA-GG splice sites. RNA-Seq reads supported 224 of these CA-GG splice sites. Flanking sequences of CA-GG and GC-AG splice sites were extracted and aligned to investigate the reason for these erroneous transcript alignments (Additional file 13). An additional G directly downstream of the 3’ AG splice site was only present when this splice site was predicted as GG. Cases where the GC-AG was predicted lack this G thus preventing the annotation of a CA-GG splice site combination.

### Non-canonical splice sites in single copy genes

To assess the impact of gene copy number on the presence of non-canonical splice sites, we compared a group of presumably single copy genes against all other genes. The average percentage of genes with non-canonical splice sites among single copy BUSCO genes was 11.4%. The average percentage among all genes was only 10.4%. This uncorrected difference between both groups is statistically significant (p=0.04, Mann-Whitney U test), but species-specific effects were obvious. While the percentage in some species is almost the same, other species show a much higher percentage of genes with non-canonical splice sites among BUSCO genes (Additional file 14). A couple of species displayed an inverted situation, having less genes with non-canonical splice sites among the BUSCO genes than the genome-wide average.

### Intron analysis

Length distributions of introns with canonical and non-canonical splice site combinations are similar in most regions (Fig.3). However, there are three striking differences between both distributions: i) the higher abundance of very short introns with non-canonical splice sites, ii) the lower peak at the most frequent intron length (around 200 bp), and iii) the high percentage of introns with non-canonical splice sites that are longer than 5 kb. These distributions indicate that non-canonical splice sites are more frequent in introns that deviate from the average length. Although the total number of introns with canonical splice sites longer than 5 kb is much higher, the proportion of non-canonical splice sites containing introns is on average at least twice as high as the proportion of introns with canonical splice site combinations. These differences between both distributions are significant (Wilcoxon test, p-value=0.02). Although differences in the frequency of non-canonical splice site combinations in introns longer than 5kb exist, no clear pattern of preferred motifs was detected. However, it seems that G might be underrepresented in frequent splice sit combinations in these long introns.

**Fig. 3:**
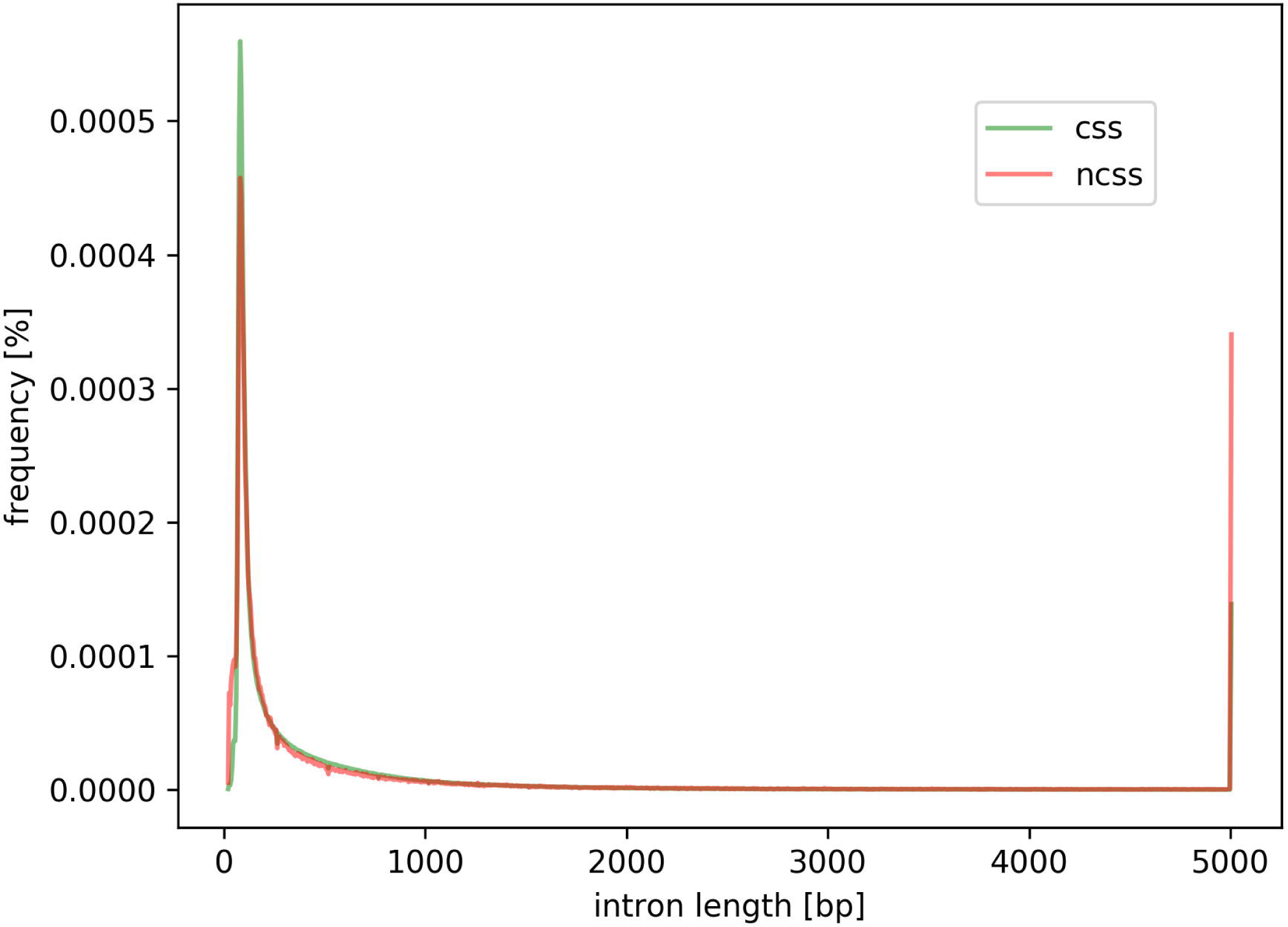
Intron length distribution. Length distribution of introns with canonical (green) and non-canonical (red) splice site combinations are displayed. Values of all species are combined in this plot resulting in a consensus curve. Most striking differences are (1) at the intron length peak around 200 bp where non-canonical splice site combinations are less likely and (2) at very long intron lengths where introns with non-canonical splice sites are more likely.

Stress-related genes were checked for increased intron sizes, because non-canonical splice site combinations might be associated with stress-response. Comparison of stress-related genes in *A. thaliana, Beta vulgaris, Brassica oleracea, B.napus, B.rapa, and Vitis vinifera* did not reveal a substantially increased intron size in these genes.

The likelihood of having a non-canonical splice site in a gene is almost perfectly correlated with the number of introns (Additional file 15). Analyzing this correlation across all plant species resulted in a sufficiently large sample size to see this effect even in genes with about 40 introns. Insufficient sample sizes kept us from investigating it for genes with even more introns.

### Conservation of non-canonical splice sites

Non-canonical splice site combinations detected in *A. thaliana* Col-0 were compared to single nucleotide polymorphisms of 1,135 accessions which were studied as part of the 1001 genomes project. Of 1,296 non-canonical splice site combinations, 109 overlapped with listed variant positions. At 21 of those positions, the majority of all accessions displayed the Col-0 allele, while the remaining 88 positions were dominated by other alleles.

To differentiate between randomly occurring non-canonical splice sites (e.g. sequencing errors) and true biological variation, the conservation of non-canonical splice sites across multiple species can be analyzed. This approach was demonstrated for the selected candidate At1g79350 (rna15125). Manual inspection revealed that non-canonical splice sites were conserved in three positions in many putative homologous genes across various species (Additional file 16).

### RNA-Seq-based validation of annotated splice sites

RNA-Seq reads of 35 different species (Additional file 2) were mapped to the respective genome sequence to allow the validation of splice sites based on changes in the read coverage depth (Additional file 3, Additional file 17). Validation ratios of all splice sites ranged from 75.5% in *Medicago truncatula* to 96.4% in *Musa acuminata*. A moderate correlation (r=0.46) between the amount of RNA-Seq reads and the ratio of validated splice sites was observed (Additional file 18). When only considering non-canonical splice sites, the validation ranged from 15.2% to 91.3% displaying a similar correlation with the amount of sequencing reads. Based on validated splice sites, the proportion of different splice site combinations was analyzed across all species (Fig.4). The average percentages are approximately 98.7% for GT-AG, 1.2% for GC-AG, 0.06% for AT-AC, and 0.09% for all other minor splice site combinations. *Medicago truncatula, Oryza sativa, Populus trichocarpa, Monoraphidium neglectum*, and *Morus notabilis* displayed substantially lower validation values for the major non-canonical splice sites.

**Fig. 4:**
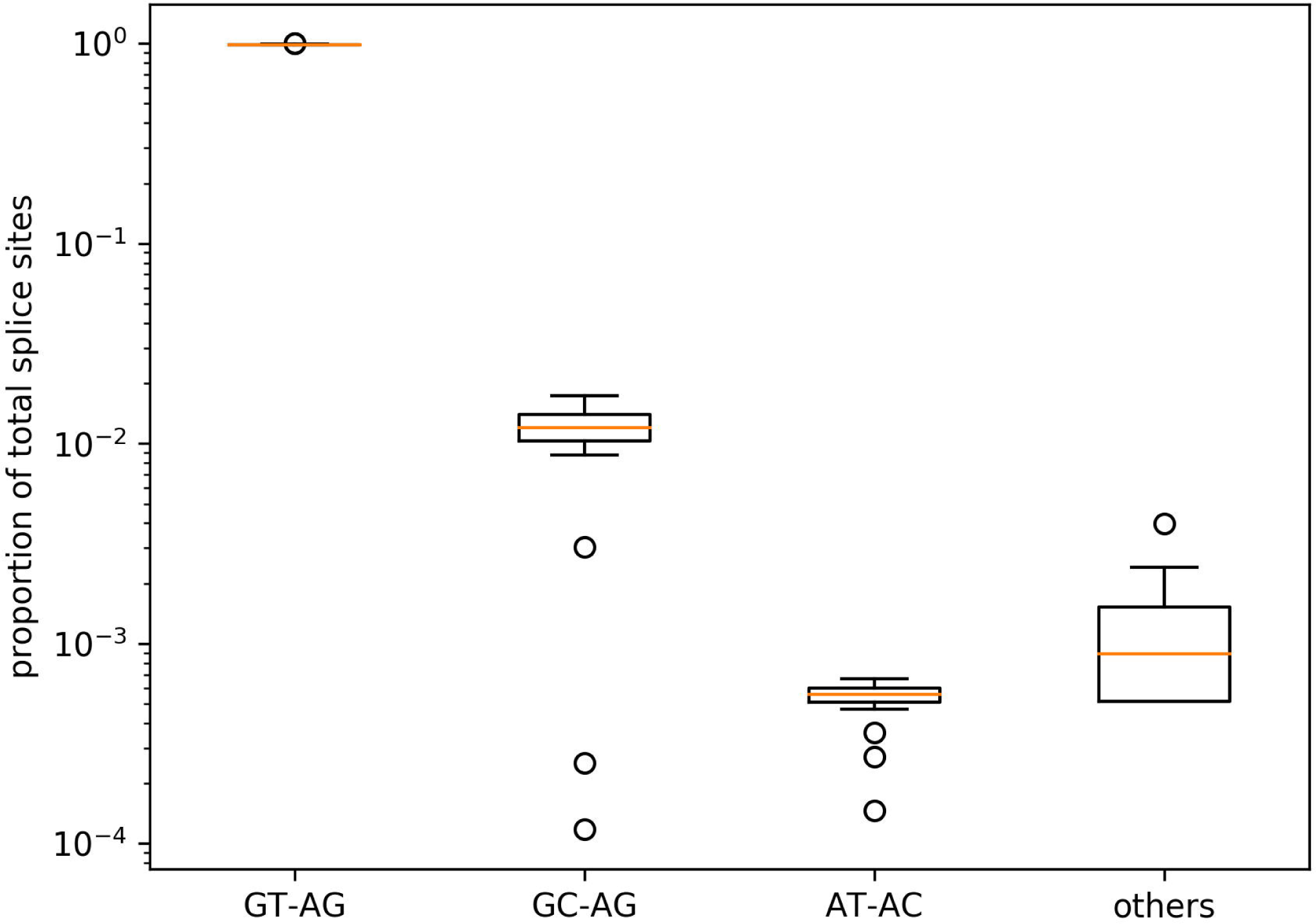
Splice site frequency. Occurrences of the canonical GT-AG, the major non-canonical GC-AG and AT-AC as well as the combined occurrences of all minor non-canonical splice sites (others) are displayed. The proportion of GT-AG is about 98.7%. There is some variation, but most species show GC-AG at about 1.2% and AT-AC at 0.06%. All others combined account usually for about 0.09% as well.

### Quantification of splice site usage

Based on mapped RNA-Seq reads, the usage of different splice sites was quantified (Fig.5; [49]). Canonical GT-AG splice site combinations displayed the strongest RNA-Seq read coverage drop when moving from an exon into an intron (Additional file 3). There was a substantial difference in average splice site usage between 5’ and the 3’ ends of GT-AG introns. The same trend holds true for major non-canonical GC-AG splice site combinations, while the total splice site usage is lower. Major non-canonical AT-AC and minor non-canonical splice sites did not show a difference between 5’ and 3’ end. However, the total usage values of AT-AC are even lower than the values of GC-AG splice sites.

**Fig. 5:**
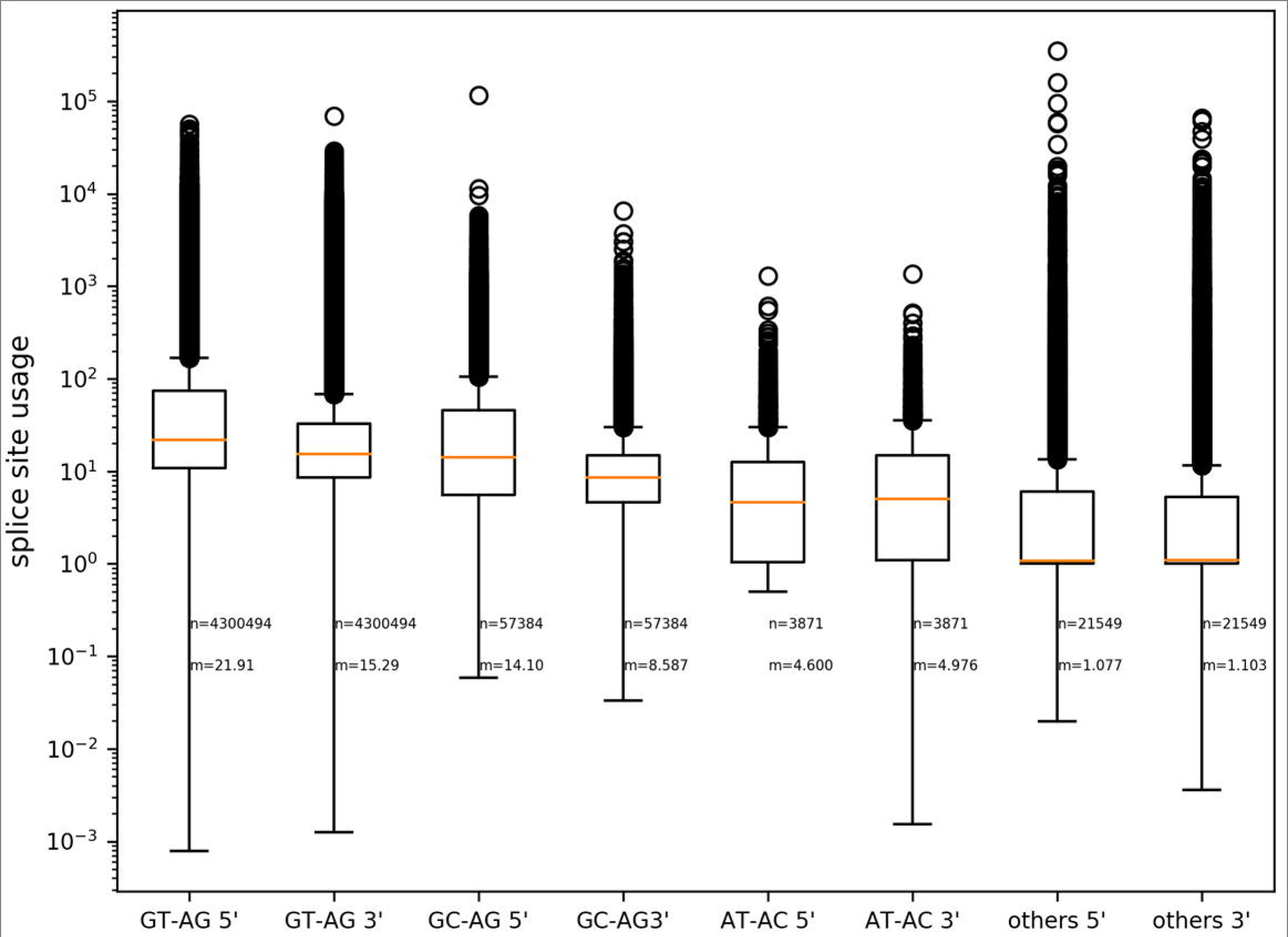
Usage of splice sites. Usage of splice sites was calculated based on the number of RNA-Seq reads supporting the exon next to a splice site and the number of reads supporting the intron containing the splice site. There is a substantial difference between the usage of 5’ and 3’ splice sites in favor of the 5’ splice sites. Canonical GT-AG splice site combinations are used more often than major or minor non-canonical splice site combinations. Sample size (n) and median (m) of the usage values are given for all splice sites.

There is a significant correlation between the usage of a 5’ splice site and the corresponding 3’ splice site. However, the Spearman correlation coefficient varies between all four groups of splice sites ranging from 0.42 in minor non-canonical splice site combinations to 0.82 in major non-canonical AT-AC splice site combinations.

In order to provide an example for the usage of minor non-canonical splice sites under stress conditions, four single RNA-Seq datasets of *B. vulgaris* were processed separately. They are the comparison of control vs. salt and control vs. high light [50]. The number of RNA-Seq supported minor non-canonical splice sites increased between control and stress conditions from 17 to 19 and from 21 to 24, respectively. GT-TA and AA-TA were only supported by RNA-Seq reads derived from samples under stress conditions.

## Discussion

This inspection of non-canonical splice sites annotated in plant genome sequences was performed to capture the diversity and to assess the validity of these annotations, because previous studies indicate that annotations of non-canonical splice sites are a mixture of artifacts and *bona fide* splice sites [29, 34, 51]. Our results update and expand previous systematic analyses of non-canonical splice sites in smaller data sets [29, 30, 33, 34]. An extended knowledge about non-canonical splice sites in plants could benefit gene predictions [30, 52], as novel genome sequences are often annotated by lifting an existing annotation.

### Confirmation of *bona fide* splicing from minor non-canonical combinations

Our analyses supported a variety of different non-canonical splice sites matching previous reports of *bona fide* non-canonical splice sites [29, 30, 34, 51]. Frequencies of different minor non-canonical splice site combinations are not random and vary between different combinations. Those combinations similar to the canonical combination or the major non-canonical splice site combinations are more frequent. Furthermore, our RNA-Seq analyses demonstrate the actual use of non-canonical splice sites, revealing a huge variety of different transcripts derived from non-canonical splice sites, which may be evolutionarily significant. Although, some non-canonical splice sites may be located in pseudogenes, the transcriptional activity and accurate splicing at most non-canonical splice sites indicates functional relevance e.g. by contributing to functional diversity as previously postulated [2, 25, 26]. These findings are consistent with published reports that have demonstrated functional RNAs generated from non-canonical splice sites [30, 53].

In general, the pattern of non-canonical splice sites is very similar between species with major non-canonical splice sites accounting for most cases of non-canonical splicing. While the average across plants of 98.7% GT-AG canonical splice sites is in agreement with recent reports for *A. thaliana* [30], it is slightly lower than 99.2 % predicted for mammals [33] or 99.3% as previously reported for Arabidopsis based on cDNAs [54]. In contrast, the frequency of major non-canonical GC-AG splice sites in plants is almost twice the value reported for mammals [33]. Most importantly the proportion of 0.09% minor non-canonical splice site combinations in plants is substantially higher than the estimation of 0.02% initially reported for mammals [33]. Taking these findings together, both major and minor non-canonical splice sites could be a more significant phenomenon of splicing in plants than in animals. This hypothesis would be consistent with the notion that splicing in plants is a more complex and diverse process than that occurring in metazoan lineages [55–57]. An in-depth investigation of non-canonical splice sites in animals and fungi would be needed to validate this hypothesis.

### Species-specific differences in minor non-canonical splice site combinations

As previous studies on non-canonical splice sites were often focused on one species [54] or a few model organisms [33, 34, 38], the observed variation among the plant genomes investigated here updates the current knowledge and revealed potential species-specific differences. However, small numbers of non-canonical splice sits in some species might prevent the detection of phylogenetic patterns in the genome-wide analysis. Nevertheless, conserved non-canonical splice site positions exist as presented on the gene level for At1g79350. Differences in the availability of hints in the gene prediction process and variation in the assembly quality might contribute to the observed differences in the number of non-canonical splice sites between closely related species.

The group of minor non-canonical splice sites displayed the largest variation between species, and a frequent non-canonical splice site combination (CA-GG) which appeared peculiar to *O. sativa* is probably due to an alignment error. In other words, the predicted CA-GG splice site combinations in rice can be conceived as major non-canonical GC-AG events by just splitting the transcript sequence in a different way during the alignment over the intron. An additional downstream G at the 3’ splice site seems to be responsible for leading to this annotation, because cases where GC-AG was correctly annotated do not display this G in the respective position. Dedicated alignment tools are needed to bioinformatically distinguish these events [58], otherwise manual inspection must be used to correctly resolve these situations.

Despite all artifacts described here and elsewhere [29, 33, 59], non-canonical splice sites seem to have conserved functions as indicated by conservation over long evolutionary periods displayed as presence in homologous sequences in multiple species [23, 29]. Our own analyses across multiple accessions of *A. thaliana* support this conjecture and suggest that some non-canonical splice sites are conserved in homologous loci at the intra-specific level. At the same time, there is intra-specific variability [30] that might be attributed to the accumulation of mutations prior to purifying selection. Assessing the variability within a species could be an additional approach to distinguish *bona fide* splice sites from artifacts or recent mutations.

### Putative mechanisms for processing of minor non-canonical splice sites

We sought to understand possible correlations with minor non-canonical splice site combinations in order understand the mechanisms driving their occurrence. Therefore, we explored the impact of genomic position relative to centromeres, the effect of increased gene number, and the impact of intron length. The occurrence of non-canonical splice sites is reduced with proximity to the centromere, but this is likely due to reduced gene content in centromeric regions. Averaged across all species, there a significantly higher proportion of non-canonical sites in single copy genes, but species-specific differences also violate this observation, suggesting that gene copy number is not an important determinant. However, non-canonical splice sites may be more important in splicing very long introns, because they appear in introns above 5 kb with a higher relative likelihood than canonical splice sites. Further investigations are needed to validate the observed lack of G in these splice site combinations and to identify an underlying pattern if it exists. When looking for an association of long introns with stress-related genes, no significant increase in their intron sizes was observed. However, it is still possible that these long introns belong to genes which were not previously described in relation to stress.

Previous studies postulated different non-spliceosomal removal mechanisms for such introns including the IRE1 / tRNA ligase system [60, 61] and short direct repeats leading to transcriptional slippage [62, 63]. It should be mentioned that many sequence variants of snRNAs are encoded in plant genomes [64]. The presence of multiple spliceosome types in addition to the canonical U2 and the non-canonical U12 spliceosome could be another explanation [38].

Another hypothesis suggests parasitic splice sites using neighbouring recognition sites for the splicing machinery to enable their processing [33]. The mere presence of GT close to the 5’ non-canonical splice site and AG close to the 3’ non-canonical splice site might be sufficient for this process to take place. These non-canonical splice sites are expected to be in frame with the associated GT-AG signals which could be responsible for recruiting the splicing machinery [33]. This hypothesis is supported by the observation that splice sites seem to be missed sometimes thus leading to the use of the next splice site which is usually in frame with the original one [54]. Further investigation might connect neighbouring sequences to the processing of minor non-canonical splice sites.

There is no evidence for RNA editing to modify splice sites yet, but previous studies found that modifications of mRNAs are necessary to enable proper splicing in some cases [65]. Even so such a system is probably not in place for all minor non-canonical splice sites, a modification of nucleotides in the transcript would be another way to regulate gene expression at the post-transcriptional level.

Although, these hypotheses could be an additional or alternative explanations for the situation observed in *O. sativa*, considering the CA-GG cases as annotation and alignment errors seems more likely due to their unique presence in this species.

### Usage of non-canonical splice sites

Our results could provide a strong foundation to further analyses of the splicing process by providing detailed information about the frequency at which splicing occurred at a certain splice site. The results indicate that this usage of different splice site types could vary substantially. A possible explanation for these observed differences is the mixture of RNA-Seq data sets, which contains samples from various tissues and different environmental or physiological conditions. Sequencing reads reflect the splicing events occurring under these specific conditions. As previously indicated by several reports, non-canonical splice sites might be more frequently used under stress conditions [25, 51, 63]. As most plants are unable to escape environmental conditions by movement, a higher frequency of non-canonical splice sites in sessile plant species compared to other taxonomic groups should be assessed in the future to explore whether there may be a link between non-canonical splice frequency and life habit.

The observation of a stronger usage of the donor splice site over the acceptor splice site in GT-AG and GC-AG splice site combinations is matching previous reports where one donor splice site can be associated with multiple acceptor splice sites [54, 66]. The absence of this effect at minor non-canonical splice site combinations might hint towards a different splicing mechanism, which is restricted to precisely one combination of donor and acceptor splice site.

The observed usage of GT-TA and AA-TA splice site combinations under stress conditions in contrast to control conditions as well as the slight increase in the number of supported minor non-canonical splice site combinations requires further testing e.g. in other species or under different stress conditions. It would be interesting to validate the usage of different splice sites in response to stress and not just the expression of stress-related genes. In principle, it would be possible to assess the usage of splice sites under diverse environmental or developmental conditions as performed in this study for different plant species. While numerous RNA-Seq datasets are available per species, these analyses would require a large number of datasets generated under identical or at least similar conditions. Therefore, the identification of splicing variants dedicated to certain stress responses is beyond the scope of this work.

### Limitations of the current analyses

Some constraints limit the power of the presented analyses. In accordance with the important plant database Araport11 [37] and previous analyses [30], only the transcript encoding the longest peptide sequence was considered when investigating splice site conservation across species. Although the exclusion of alternative transcripts was necessary to compensate differences in the annotation quality, more non-canonical splice sites could be revealed by investigations of all transcript versions in the future. The exclusion of annotated introns shorter than 20 bp as well as the minimal intron length cutoff of 20 bp during the RNA-Seq read mapping prevented the investigation of very small introns. There are reports of experimentally validated introns with a minimal length of 56bp [67]. Although recent reports indicate a minimal intron length around30 bp in humans [68] or even down to 10 bp [51], it is unclear if very short sequences should be called introns. Since spliceosomal removal of these very short sequences via lariat formation seems unlikely, a new terminology might be needed. The applied length cutoff was selected to avoid previously reported issues with false positives [51]. However, d*e novo* identification of very short introns as recently performed for *Mus musculus* and *H. sapiens* [51, 69] could become feasible as RNA-Seq data sets based on similar protocols become available for a broad range of plant species. Variations between RNA-Seq samples posed another challenge. Since there is a substantial amount of variation within species [70, 71], we can assume that small differences in the genetic background of the analyzed material could bias the results. Splice sites of interest might be canonical splice site combinations in some accessions or subspecies, respectively, while they are non-canonical in others. Despite our attempts to collect RNA-Seq samples derived from a broad range of different conditions and tissues for each species, data of many specific physiological states are missing for most species. Therefore, we cannot exclude that certain non-canonical splice sites were missed in our splice site usage analysis due to a lack of gene expression under the investigated conditions.

### Future Perspectives

As costs for RNA-Seq data generation drops over the years [72], improved analyses will become possible over time. Investigation of homologous non-canonical splice sites poses several difficulties, as the exonic sequence is not necessarily conserved. Due to upstream changes in the exon-intron structure [73], the number of the non-canonical introns can differ between species. However, a computationally feasible approach to investigate the phylogeny of all non-canonical splice sites would significantly enhance our knowledge e.g. about the emergence and loss of non-canonical splice sites. Experimental validation of splice sites *in vivo* and *in vitro* could be the next step. It is crucial for such analyses to avoid biases introduced by reverse transcription artifacts e.g. by comparing different enzymes and avoiding random hexameters during cDNA synthesis [74]. Splice sites could be experimentally validated e.g. by integration in the *Aequoria vicotria* GFP sequence [75] to see if they are functional in plants. Our analyses support the concept that differences between plant species need to be taken into account when performing such investigations [76, 77].

## Supporting information

## Declarations

### Ethics approval and consent to participate

Not applicable

### Consent for publication

Not applicable

### Availability of data and materials

The datasets generated during the current study are included as Additional files and publicly available from https://doi.org/10.4119/unibi/2931315. Scripts written for the described analyses are available on github: https://github.com/bpucker/ncss2018.

### Authors’ contribution

BP and SFB designed the research. BP performed bioinformatic analyses. BP and SFB interpreted the results and wrote the manuscript.

## Acknowledgements

We are thankful to everyone involved in generating the datasets underlying this study.

## Additional files

**Additional file 1. Analysed data sets.** List of investigated genome sequences and corresponding annotation. Md5sums are given for all files.

**Additional file 2. RNA-Seq data sets.** List of Sequence Read Archive accession numbers of all included RNA-Seq data sets sorted by species.

**Additional file 3. RNA-Seq based splice site validation.** Schematic illustration how the splitted mapping of RNA-Seq reads (arrows) over exons (red) and introns (grey) was used to validate splice sites. The read coverage depth should drop when moving from an exon into an intron. Red arrows indicate the four positions considered for this analysis.

**Additional file 4. Phylogenetic tree.** RbcL sequences were used to construct a phylogenetic tree of all species involved in the analysis. Missing data points were corrected by relying on the NCBI taxonomy thus the branch lengths are not to scale.

**Additional file 5. Genome statistics.** Statistical information about each analyzed genome sequence and the average values across all species are listed.

**Additional file 6. Number of splice sites per species.** Canonical and non-canonical splice sites were counted per species as described in the method section.

**Additional file 7. Splice site diversity per species.** The occurrence of all possible splice site combinations was counted for all species as described in the method section.

**Additional file 8. Similarity of the non-canonical splice site pattern across plants.** The Spearman correlation coefficient between each pair of plants was calculated based on the observed frequency of all possible splice site combinations. Red color indicates similarity while blue color indicates substantial differences. As this correlation calculation takes the individual counts for all splice sites combinations in all species into account, it is possible to calculate correlation values even in the absence of non-canonical splice sites.

**Additional file 9. Correlation of splice site frequencies with genome size.** For each investigated species the number of canonical and non-canonical splice sites is displayed. The Spearman correlation coefficient between splice site number and genome size is r=0.14 for canonical splice sites and r=0.02 for non-canonical splice sites.

**Additional file 10. Genome-wide distribution of non-canonical splice sites in *A. thaliana***. The distribution of genes with non-canonical splice sites (red dots) across the five chromosome sequences (black lines) of *A. thaliana* was analysed.

**Additional file 11. Genome-wide distribution of non-canonical splice sites in *B. vulgaris***. The distribution of genes with non-canonical splice sites (red dots) across the nine chromosome sequences (black lines) of *B. vulgaris* was analysed.

**Additional file 12. Genome-wide distribution of non-canonical splice sites in *V.vinifera***. The distribution of genes with non-canonical splice sites (red dots) across the nineteen chromosome sequences (black lines) of *V. vinifera* was analysed.

**Additional file 13. Conserved sequences around splice sites in *Oryza sativa***. Predicted splice site combinations observed in *Oryza sativa* are indicated by a black line below them. Donor splice sites are on the left, acceptor splice sites on the right. The minor non-canonical splice combination CA-GG at the top could be converted into the major non-canonical GC-AG combination by just shifting one nucleotide to the left. The presence of two Gs at the acceptor splice site seems to correlate with the prediction of this CA-GG splice site combination instead of a major non-canonical GC-AG.

**Additional file 14. Non-canonical splice sites in single copy genes.** The occurrence of non-canonical splice sites in single copy genes (BUSCO) and in all genes was assessed per species.

**Additional file 15. Proportion of non-canonical splice sites.** The green line indicates the average (median) proportion of genes with a non-canonical splice site combination. Grey lines indicate the range between 25% and 75% quantiles. Genes with more introns are more likely to have a non-canonical splice site combination. There is an almost perfect correlation up to 40 introns per gene. Insufficient sample sizes above this intron number prevent further analyses.

**Additional file 16. Conservation of non-canonical splice sites.** Non-canonical splice sites at conserved positions in putative homologous of At1g79350 across various species.

**Additional file 17. Supported splice sites.** Percentage of splice sites supported by RNA-Seq reads is given per species.

**Additional file 18. RNA-Seq data set sizes.** There is a moderate correlation between the amount of bases in the used RNA-Seq data sets and the number of supported splice sites. The trend is similar for canonical (r=0.46) and non-canonical (r=0.43) splice site combinations.

## References

1. Berget SM, Moore C, Sharp PA. Spliced segments at the 5’ terminus of adenovirus 2 late mRNA. Proc Natl Acad Sci U S A. 1977;74:3171–5.

2. Gilbert W. The Exon Theory of Genes. Cold Spring Harb Symp Quant Biol. 1987;52:901–5.

3. Koonin EV, Senkevich TG, Dolja VV. The ancient Virus World and evolution of cells. Biol Direct. 2006;1:29.

4. Carmel L, Chorev M. The Function of Introns. Front Genet. 2012;3. doi:https://doi.org/10.3389/fgene.2012.00055.

5. Jo B-S, Choi SS. Introns: The Functional Benefits of Introns in Genomes. Genomics Inform. 2015;13:112–8.

6. Mukherjee D, Saha D, Acharya D, Mukherjee A, Chakraborty S, Ghosh TC. The role of introns in the conservation of the metabolic genes of Arabidopsis thaliana. Genomics. 2018;110:310–7.

7. Rogozin IB, Carmel L, Csuros M, Koonin EV. Origin and evolution of spliceosomal introns. Biol Direct. 2012;7:11.

8. Csuros M, Rogozin IB, Koonin EV. A Detailed History of Intron-rich Eukaryotic Ancestors Inferred from a Global Survey of 100 Complete Genomes. PLoS Comput Biol. 2011;7. doi:10.1371/journal.pcbi.1002150.

9. Worden AZ, Lee J-H, Mock T, Rouzé P, Simmons MP, Aerts AL, et al. Green evolution and dynamic adaptations revealed by genomes of the marine picoeukaryotes Micromonas. Science. 2009;324:268–72.

10. Huff JT, Zilberman D, Roy SW. Mechanism for DNA transposons to generate introns on genomic scales. Nature. 2016;538:533–6.

11. Zimmerly S, Semper C. Evolution of group II introns. Mob DNA. 2015;6. doi:10.1186/s13100-015-0037-5.

12. Knoop V, Brennicke A. Promiscuous mitochondrial group II intron sequences in plant nuclear genomes. J Mol Evol. 1994;39:144–50.

13. Pucker B, Holtgraewe D, Stadermann KB, Frey K, Huettel B, Reinhardt R, et al. A Chromosome-level Sequence Assembly Reveals the Structure of the Arabidopsis thaliana Nd-1 Genome and its Gene Set. bioRxiv 407627. doi:https://doi.org/10.1101/407627.

14. Wahl MC, Will CL, Lührmann R. The Spliceosome: Design Principles of a Dynamic RNP Machine. Cell. 2009;136:701–18.

15. Papasaikas P, Valcárcel J. The Spliceosome: The Ultimate RNA Chaperone and Sculptor. Trends Biochem Sci. 2016;41:33–45.

16. Turunen JJ, Niemelä EH, Verma B, Frilander MJ. The significant other: splicing by the minor spliceosome. Wiley Interdiscip Rev RNA. 2013;4:61–76.

17. Hall SL, Padgett RA. Conserved Sequences in a Class of Rare Eukaryotic Nuclear Introns with Nonconsensus Splice Sites. J Mol Biol. 1994;239:357–65.

18. Wu Q, Krainer AR. Splicing of a divergent subclass of AT-AC introns requires the major spliceosomal snRNAs. RNA N Y N. 1997;3:586–601.

19. Dietrich RC, Incorvaia R, Padgett RA. Terminal intron dinucleotide sequences do not distinguish between U2- and U12-dependent introns. Mol Cell. 1997;1:151–60.

20. Lewandowska D, Simpson CG, Clark GP, Jennings NS, Barciszewska-Pacak M, Lin C-F, et al. Determinants of Plant U12-Dependent Intron Splicing Efficiency. Plant Cell. 2004;16:1340–52.

21. Wang G-S, Cooper TA. Splicing in disease: disruption of the splicing code and the decoding machinery. Nat Rev Genet. 2007;8:749–61.

22. Will CL, Lührmann R. Spliceosome Structure and Function. Cold Spring Harb Perspect Biol. 2011;3:a003707.

23. Rogozin IB, Wolf YI, Sorokin AV, Mirkin BG, Koonin EV. Remarkable Interkingdom Conservation of Intron Positions and Massive, Lineage-Specific Intron Loss and Gain in Eukaryotic Evolution. Curr Biol. 2003;13:1512–7.

24. Jacob M, Gallinaro H. The 5’ splice site: phylogenetic evolution and variable geometry of association with U1RNA. Nucleic Acids Res. 1989;17:2159–80.

25. Sibley CR, Blazquez L, Ule J. Lessons from non-canonical splicing. Nat Rev Genet. 2016;17:407–21.

26. Gorlova O, Fedorov A, Logothetis C, Amos C, Gorlov I. Genes with a large intronic burden show greater evolutionary conservation on the protein level. BMC Evol Biol. 2014;14:50.

27. Sibley CR, Emmett W, Blazquez L, Faro A, Haberman N, Briese M, et al. Recursive splicing in long vertebrate genes. Nature. 2015;521:371–5.

28. Zhao W, Cheng Y, Zhang C, You Q, Shen X, Guo W, et al. Genome-wide identification and characterization of circular RNAs by high throughput sequencing in soybean. Sci Rep. 2017;7:5636.

29. Jackson IJ. A reappraisal of non-consensus mRNA splice sites. Nucleic Acids Res. 1991;19:3795–8.

30. Pucker B, Holtgräwe D, Weisshaar B. Consideration of non-canonical splice sites improves gene prediction on the Arabidopsis thaliana Niederzenz-1 genome sequence. BMC Res Notes. 2017;10. doi:https://doi.org/10.1186/s13104-017-2985-y.

31. Dobin A, Davis CA, Schlesinger F, Drenkow J, Zaleski C, Jha S, et al. STAR: ultrafast universal RNA-seq aligner. Bioinformatics. 2013;29:15–21.

32. Dobin A, Gingeras TR. Mapping RNA-seq Reads with STAR. Curr Protoc Bioinforma. 2015;51:11.14.1–11.14.19.

33. Burset M, Seledtsov IA, Solovyev VV. Analysis of canonical and non-canonical splice sites in mammalian genomes. Nucleic Acids Res. 2000;28:4364–75.

34. Sheth N, Roca X, Hastings ML, Roeder T, Krainer AR, Sachidanandam R. Comprehensive splice-site analysis using comparative genomics. Nucleic Acids Res. 2006;34:3955–67.

35. Souvorov A, Kapustin Y, Kiryutin B, Chetvernin V, Tatusova T, Lipman D. Gnomon – NCBI eukaryotic gene prediction tool. 2010. http://www.ncbi.nlm.nih.gov/core/assets/genome/files/Gnomon-description.pdf. Accessed 25 Sep 2018.

36. Simão FA, Waterhouse RM, Ioannidis P, Kriventseva EV, Zdobnov EM. BUSCO: assessing genome assembly and annotation completeness with single-copy orthologs. Bioinforma Oxf Engl. 2015;31:3210–2.

37. Cheng C-Y, Krishnakumar V, Chan AP, Thibaud-Nissen F, Schobel S, Town CD. Araport11: a complete reannotation of the Arabidopsis thaliana reference genome. Plant J. 2017;89:789–804.

38. Qu W, Cingolani P, Zeeberg BR, Ruden DM. A Bioinformatics-Based Alternative mRNA Splicing Code that May Explain Some Disease Mutations Is Conserved in Animals. Front Genet. 2017;8. doi:10.3389/fgene.2017.00038.

39. Hoff KJ, Stanke M. WebAUGUSTUS—a web service for training AUGUSTUS and predicting genes in eukaryotes. Nucleic Acids Res. 2013;41:W123–8.

40. Altschul SF, Gish W, Miller W, Myers EW, Lipman DJ. Basic local alignment search tool. J Mol Biol. 1990;215:403–10.

41. Hunter JD. Matplotlib: A 2D Graphics Environment. Comput Sci Eng. 2007;9:90–5. 42.

42. Leinonen R, Sugawara H, Shumway M, International Nucleotide Sequence Database Collaboration. The sequence read archive. Nucleic Acids Res. 2011;39 Database issue:D19-21.

43. Haak M, Vinke S, Keller W, Droste J, Rückert C, Kalinowski J, et al. High Quality de Novo Transcriptome Assembly of Croton tiglium. Front Mol Biosci. 2018;5. doi:https://doi.org/10.3389/fmolb.2018.00062.

44. Quinlan AR, Hall IM. BEDTools: a flexible suite of utilities for comparing genomic features. Bioinformatics. 2010;26:841–2.

45. Katoh K, Standley DM. MAFFT Multiple Sequence Alignment Software Version 7: Improvements in Performance and Usability. Mol Biol Evol. 2013;30:772–80.

46. Price MN, Dehal PS, Arkin AP. FastTree 2 – Approximately Maximum-Likelihood Trees for Large Alignments. PLoS ONE. 2010;5. doi:10.1371/journal.pone.0009490.

47. Hahn A, Kilian J, Mohrholz A, Ladwig F, Peschke F, Dautel R, et al. Plant Core Environmental Stress Response Genes Are Systemically Coordinated during Abiotic Stresses. Int J Mol Sci. 2013;14:7617–41.

48. Pucker B, Holtgräwe D, Sörensen TR, Stracke R, Viehöver P, Weisshaar B. A De Novo Genome Sequence Assembly of the Arabidopsis thaliana Accession Niederzenz-1 Displays Presence/Absence Variation and Strong Synteny. PLOS ONE. 2016;11:e0164321.

49. Pucker B. RNA-Seq read coverage depth of splice sites in plants. 2018. https://doi.org/10.4119/unibi/2931315. Accessed 11 Oct 2018.

50. Stracke R, Holtgräwe D, Schneider J, Pucker B, Sörensen TR, Weisshaar B. Genome-wide identification and characterisation of R2R3-MYB genes in sugar beet (Beta vulgaris). BMC Plant Biol. 2014;14:249.

51. Abebrese EL, Ali SH, Arnold ZR, Andrews VM, Armstrong K, Burns L, et al. Identification of human short introns. PLOS ONE. 2017;12:e0175393.

52. Sparks ME, Brendel V. Incorporation of splice site probability models for non-canonical introns improves gene structure prediction in plants. Bioinforma Oxf Engl. 2005;21 Suppl 3:iii20–30.

53. Gupta S, Wang B-B, Stryker GA, Zanetti ME, Lal SK. Two novel arginine/serine (SR) proteins in maize are differentially spliced and utilize non-canonical splice sites. Biochim Biophys Acta. 2005;1728:105–14.

54. Alexandrov NN, Troukhan ME, Brover VV, Tatarinova T, Flavell RB, Feldmann KA. Features of Arabidopsis Genes and Genome Discovered using Full-length cDNAs. Plant Mol Biol. 2006;60:69–85.

55. Ner-Gaon H, Leviatan N, Rubin E, Fluhr R. Comparative Cross-Species Alternative Splicing in Plants. Plant Physiol. 2007;144:1632–41.

56. Richardson DN, Rogers MF, Labadorf A, Ben-Hur A, Guo H, Paterson AH, et al. Comparative Analysis of Serine/Arginine-Rich Proteins across 27 Eukaryotes: Insights into Sub-Family Classification and Extent of Alternative Splicing. PLOS ONE. 2011;6:e24542.

57. Ling Y, Alshareef S, Butt H, Lozano-Juste J, Li L, Galal AA, et al. Pre-mRNA splicing repression triggers abiotic stress signaling in plants. Plant J. 2017;89:291–309.

58. Slater GS, Birney E. Automated generation of heuristics for biological sequence comparison. BMC Bioinformatics. 2005;6:31–31.

59. Parada GE, Munita R, Cerda CA, Gysling K. A comprehensive survey of non-canonical splice sites in the human transcriptome. Nucleic Acids Res. 2014;42:10564–78.

60. Sidrauski C, Cox JS, Walter P. tRNA Ligase Is Required for Regulated mRNA Splicing in the Unfolded Protein Response. Cell. 1996;87:405–13.

61. Gonzalez TN, Sidrauski C, Dörfler S, Walter P. Mechanism of non-spliceosomal mRNA splicing in the unfolded protein response pathway. EMBO J. 1999;18:3119–32.

62. Ritz K, van Schaik BDC, Jakobs ME, Aronica E, Tijssen MA, van Kampen AHC, et al. Looking ultra deep: Short identical sequences and transcriptional slippage. Genomics. 2011;98:90–5.

63. Dubrovina AS, Kiselev KV, Zhuravlev YN. The Role of Canonical and Noncanonical Pre-mRNA Splicing in Plant Stress Responses. BioMed Res Int. 2013;2013. doi:10.1155/2013/264314.

64. Solymosy F, Pollák T. Uridylate-Rich Small Nuclear RNAs (UsnRNAs), Their Genes and Pseudogenes, and UsnRNPs in Plants: Structure and Function. A Comparative Approach. Crit Rev Plant Sci. 1993;12:275–369.

65. Castandet B, Choury D, Bégu D, Jordana X, Araya A. Intron RNA editing is essential for splicing in plant mitochondria. Nucleic Acids Res. 2010;38:7112–21.

66. Mühlemann O, Kreivi JP, Akusjärvi G. Enhanced splicing of nonconsensus 3’ splice sites late during adenovirus infection. J Virol. 1995;69:7324–7.

67. Sasaki-Haraguchi N, Shimada MK, Taniguchi I, Ohno M, Mayeda A. Mechanistic insights into human pre-mRNA splicing of human ultra-short introns: Potential unusual mechanism identifies G-rich introns. Biochem Biophys Res Commun. 2012;423:289–94.

68. Piovesan A, Caracausi M, Ricci M, Strippoli P, Vitale L, Pelleri MC. Identification of minimal eukaryotic introns through GeneBase, a user-friendly tool for parsing the NCBI Gene databank. DNA Res Int J Rapid Publ Rep Genes Genomes. 2015;22:495–503.

69. Bai Y, Ji S, Wang Y. IRcall and IRclassifier: two methods for flexible detection of intron retention events from RNA-Seq data. BMC Genomics. 2015;16:S9.

70. Clark RM, Schweikert G, Toomajian C, Ossowski S, Zeller G, Shinn P, et al. Common Sequence Polymorphisms Shaping Genetic Diversity in Arabidopsis thaliana. Science. 2007;317:338–42.

71. Alonso-Blanco C, Andrade J, Becker C, Bemm F, Bergelson J, Borgwardt KM, et al. 1,135 Genomes Reveal the Global Pattern of Polymorphism in Arabidopsis thaliana. Cell. 2016;166:481–91.

72. Muir P, Li S, Lou S, Wang D, Spakowicz DJ, Salichos L, et al. The real cost of sequencing: scaling computation to keep pace with data generation. Genome Biol. 2016;17:53.

73. Garcia-España A, Mares R, Sun T-T, DeSalle R. Intron Evolution: Testing Hypotheses of Intron Evolution Using the Phylogenomics of Tetraspanins. PLoS ONE. 2009;4. doi:10.1371/journal.pone.0004680.

74. Houseley J, Tollervey D. Apparent Non-Canonical Trans-Splicing Is Generated by Reverse Transcriptase In Vitro. PLoS ONE. 2010;5. doi:10.1371/journal.pone.0012271.

75. Haseloff J, Siemering KR, Prasher DC, Hodge S. Removal of a cryptic intron and subcellular localization of green fluorescent protein are required to mark transgenic Arabidopsis plants brightly. Proc Natl Acad Sci U S A. 1997;94:2122–7.

76. Keith B, Chua N-H. Monocot and dicot pre-mRNAs are processed with different efficiencies in transgenic tobacco. EMBO J. 1986;5:2419–25.

77. Goodall GJ, Filipowicz W. Different effects of intron nucleotide composition and secondary structure on pre-mRNA splicing in monocot and dicot plants. EMBO J. 1991;10:2635–44.

