## Supplementary figures and images for "Genome-wide analyses supported by RNA-Seq reveal non-canonical splice sites in plant genomes"

### Supplementary file 3

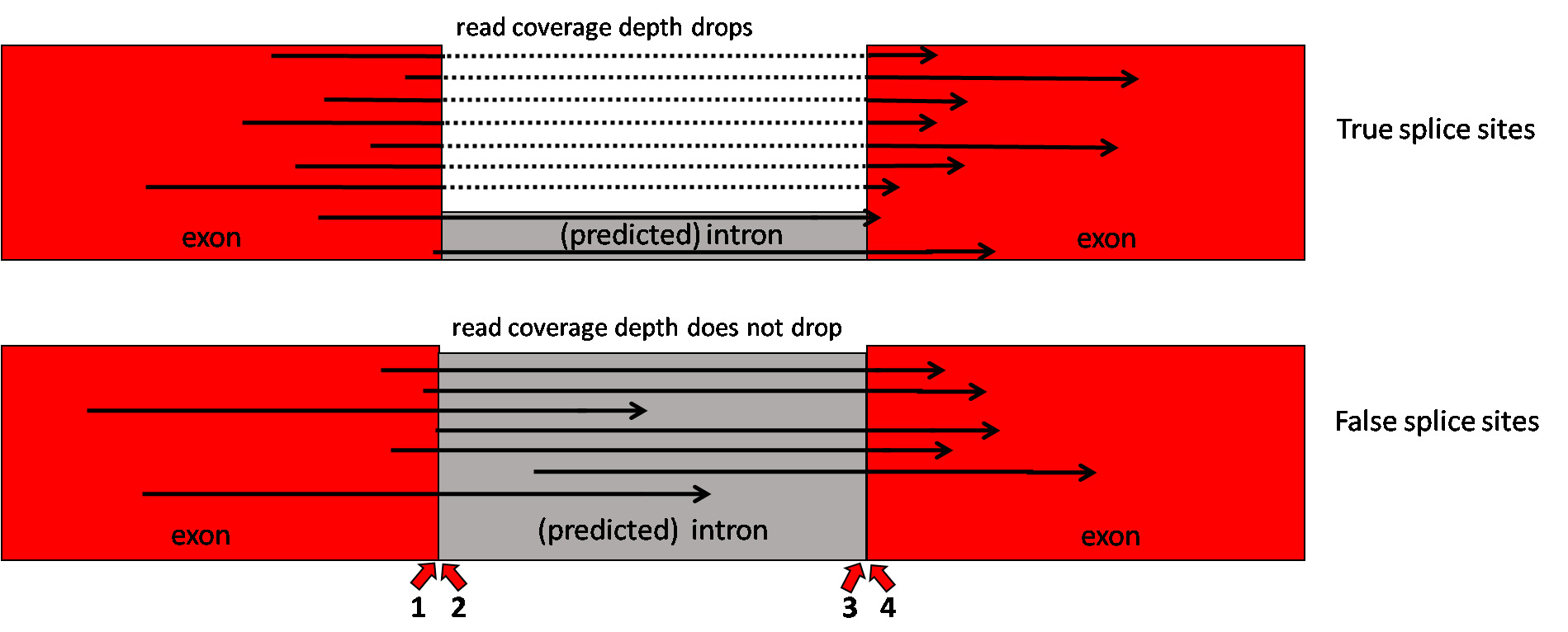

### Supplementary file 4

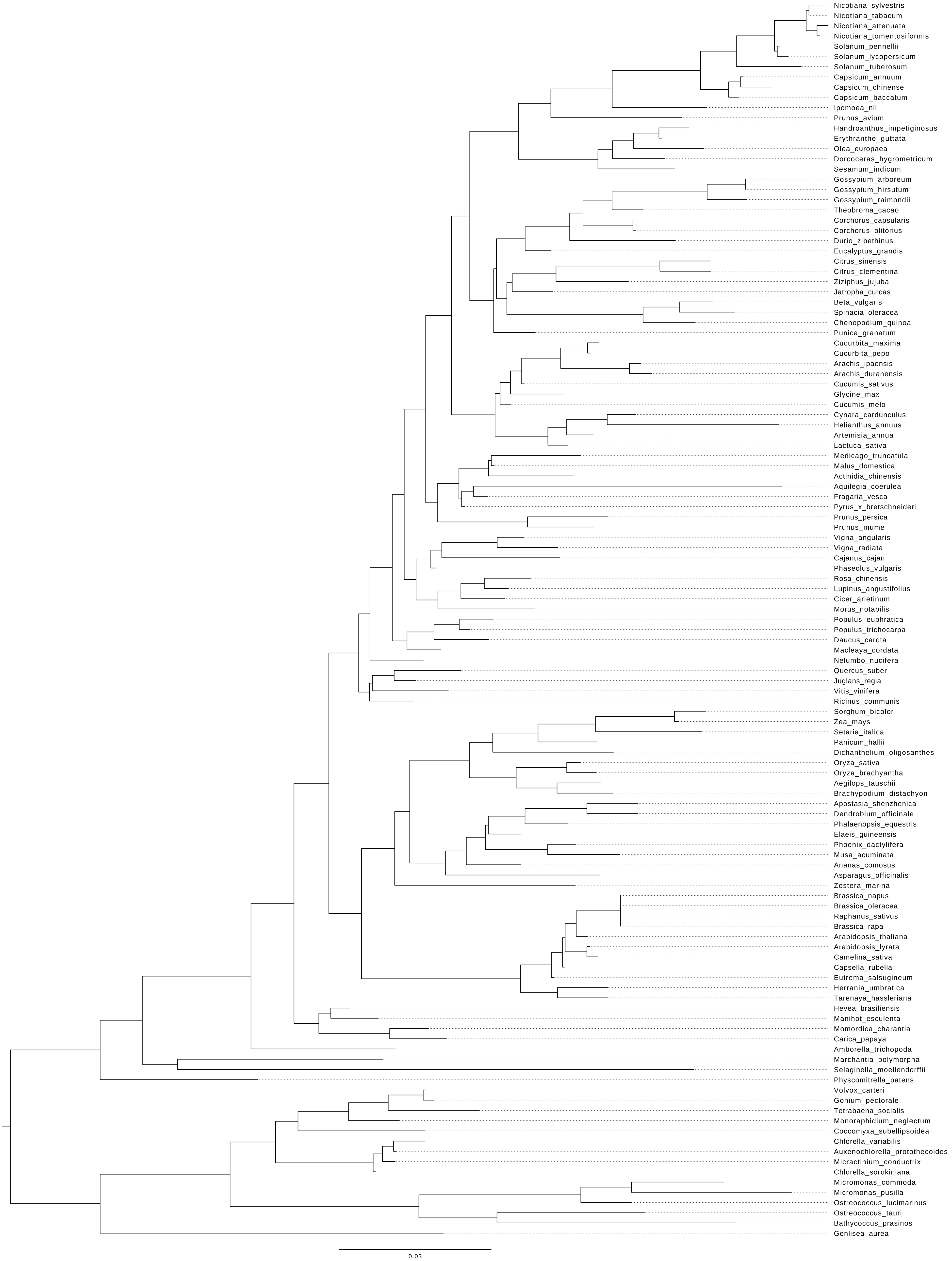

### Supplementary file 8

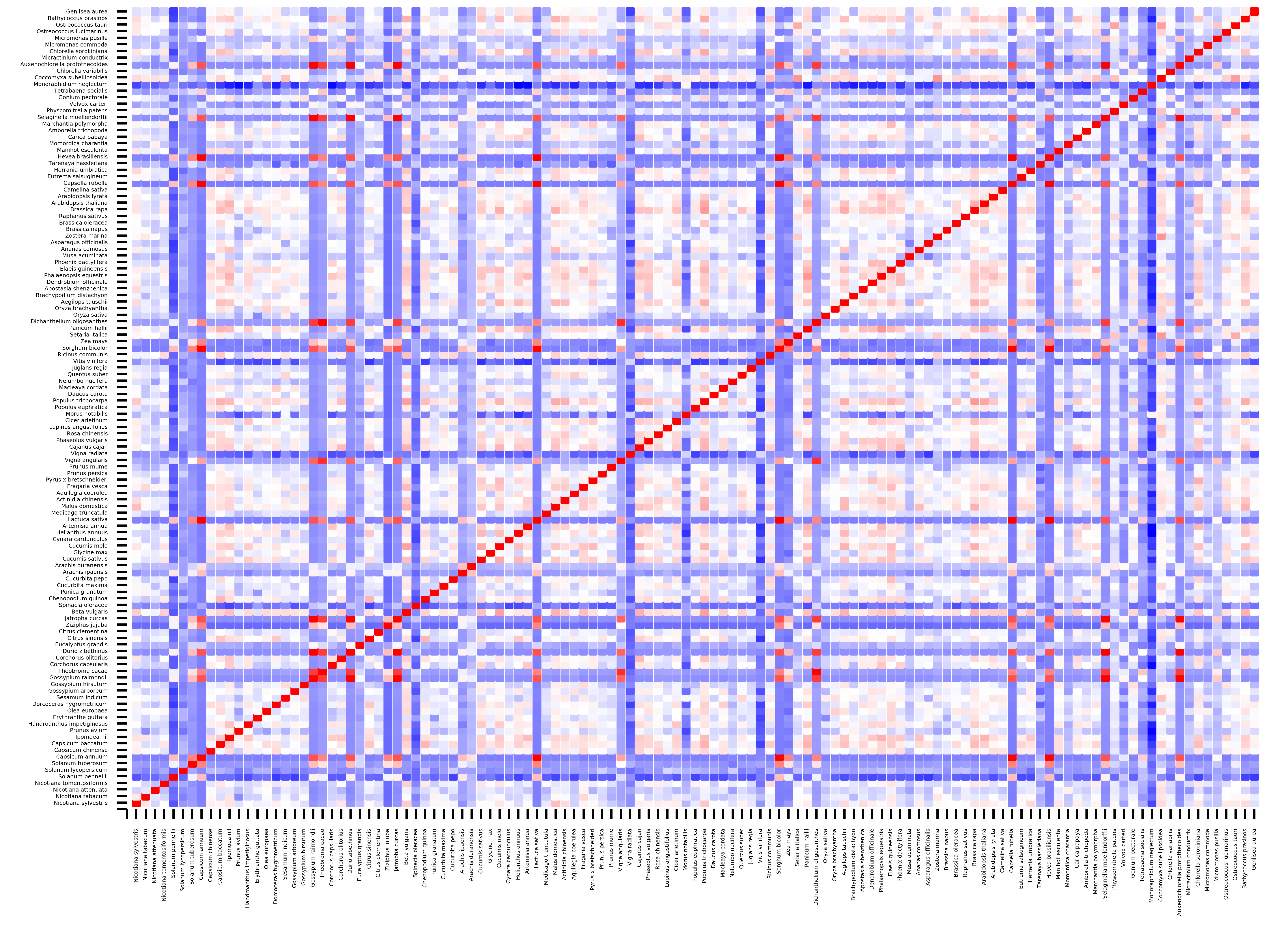

### Supplementary file 9

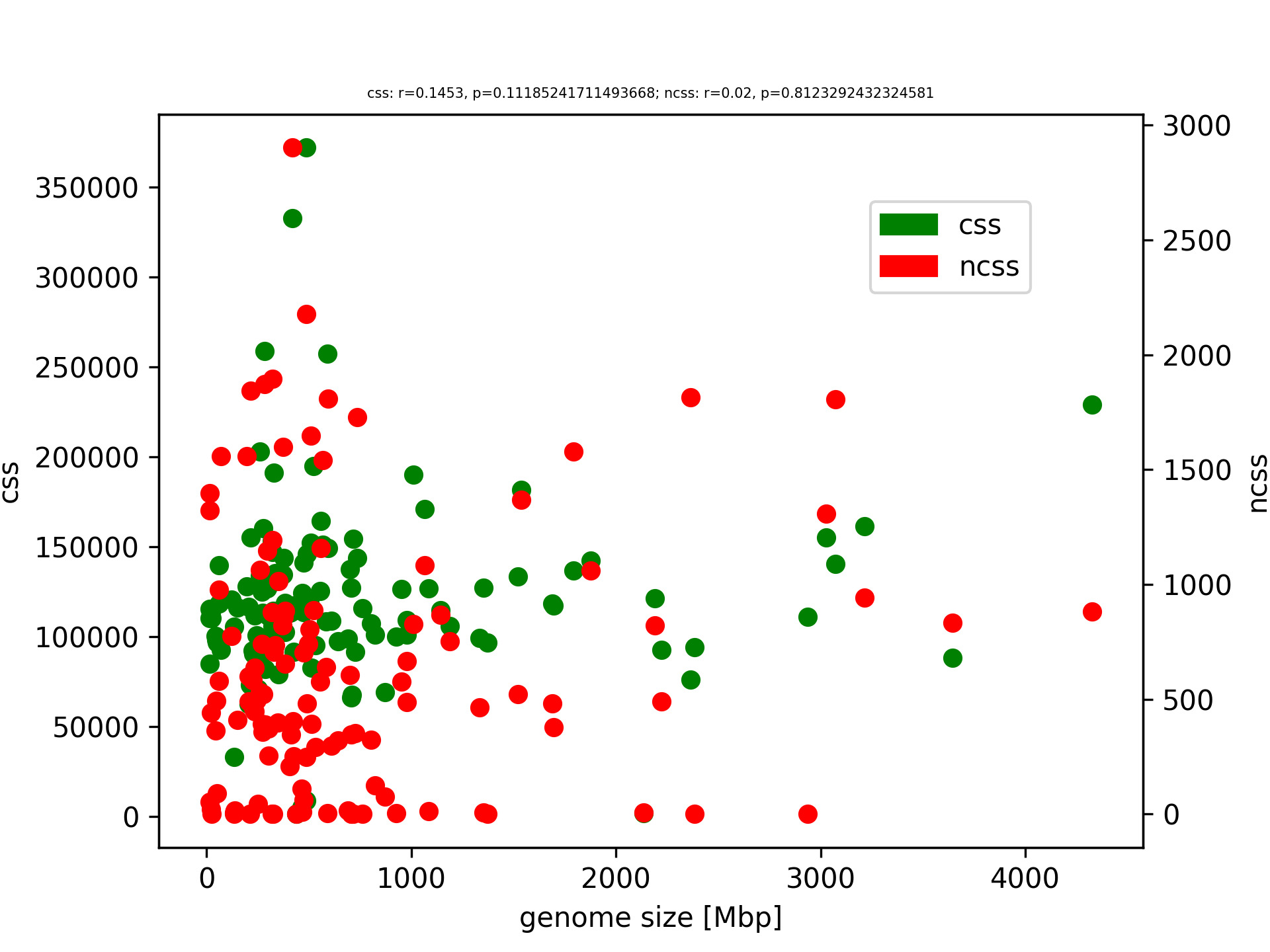

### Supplementary file 10

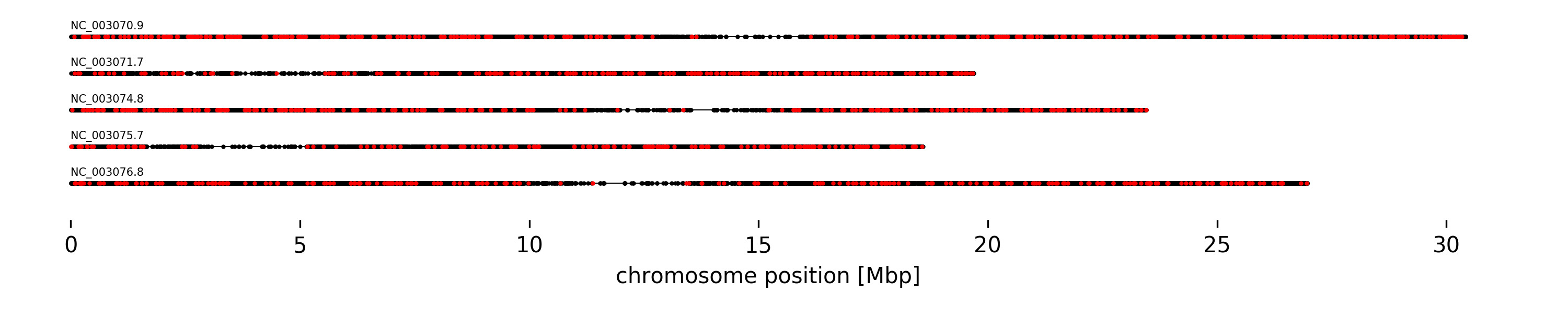

### Supplementary file 11

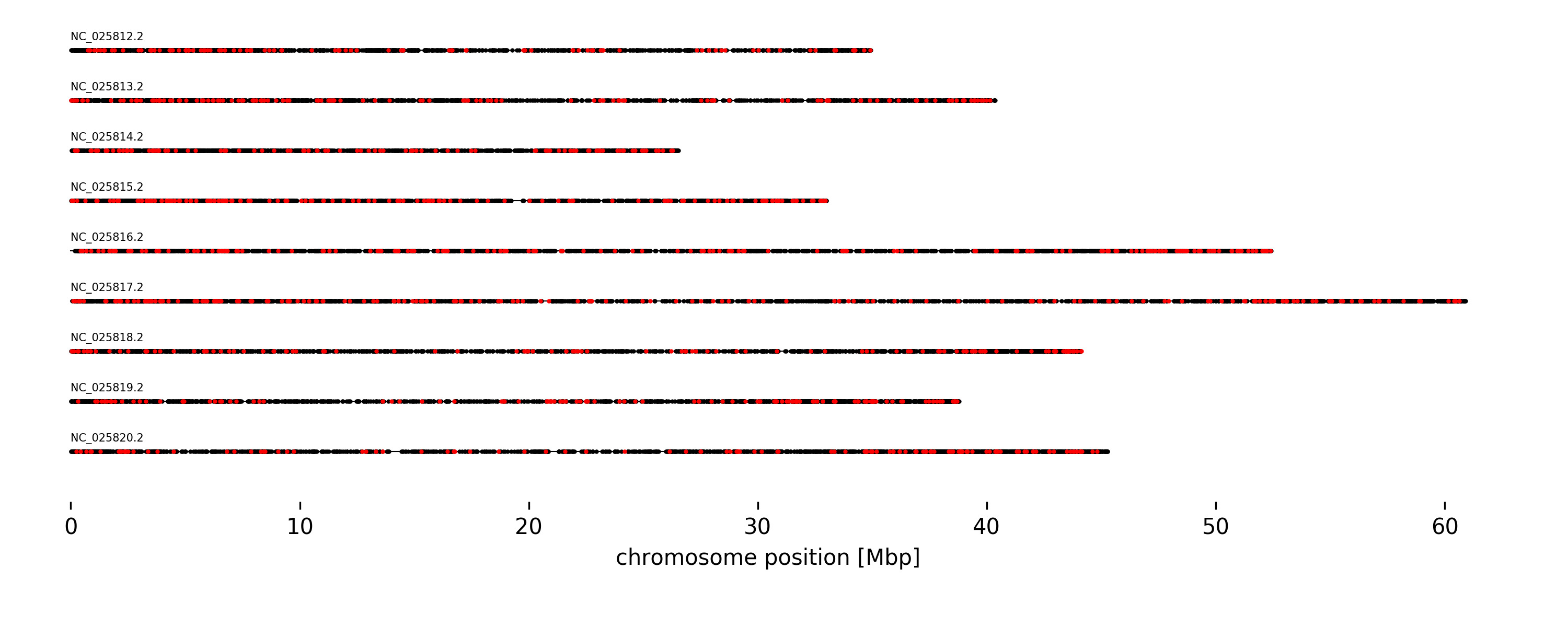

### Supplementary file 12

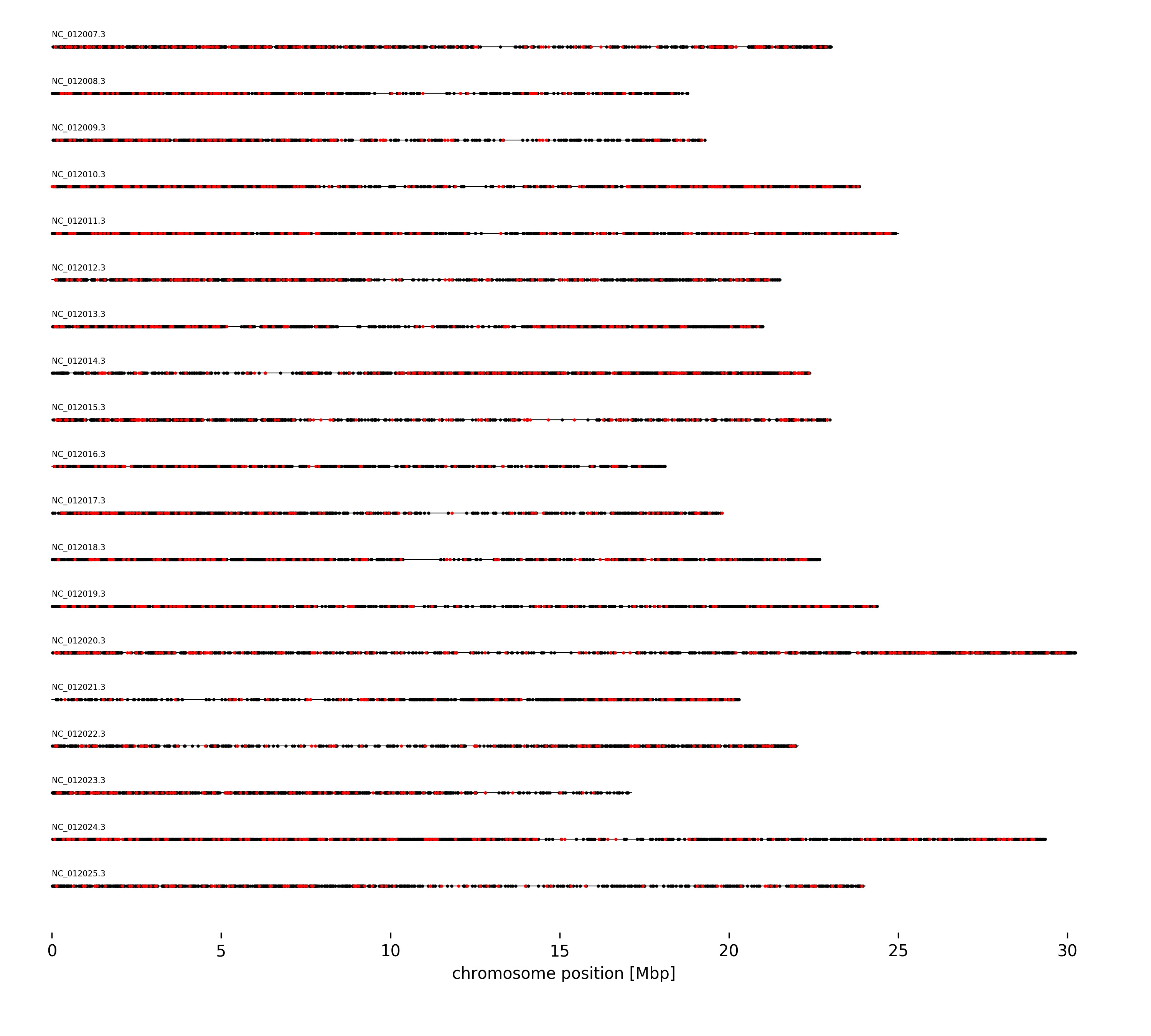

### Supplementary file 13

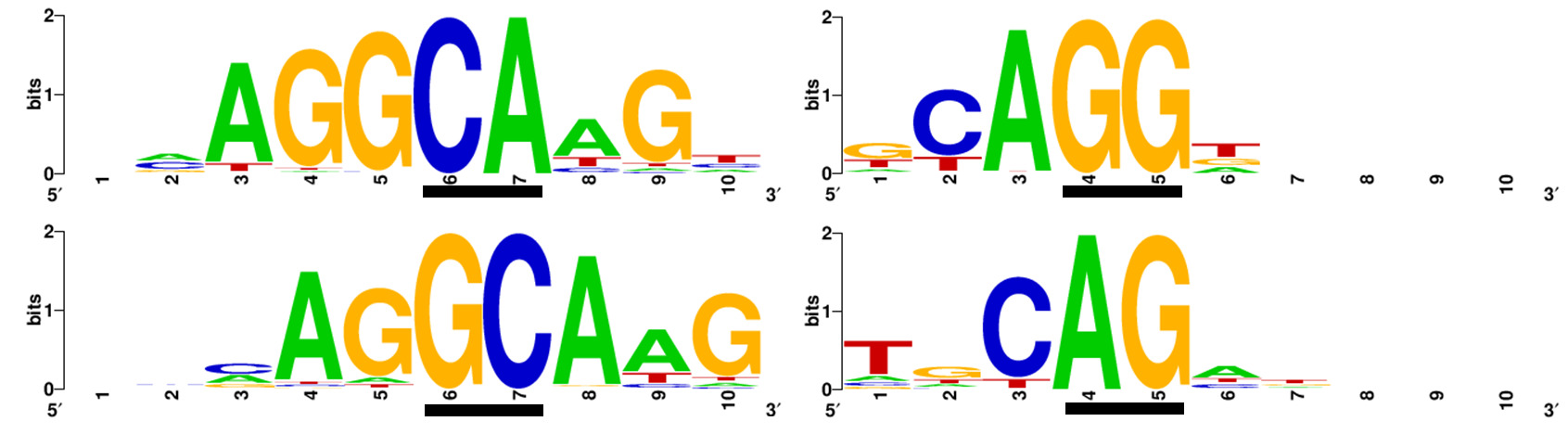

### Supplementary file 15

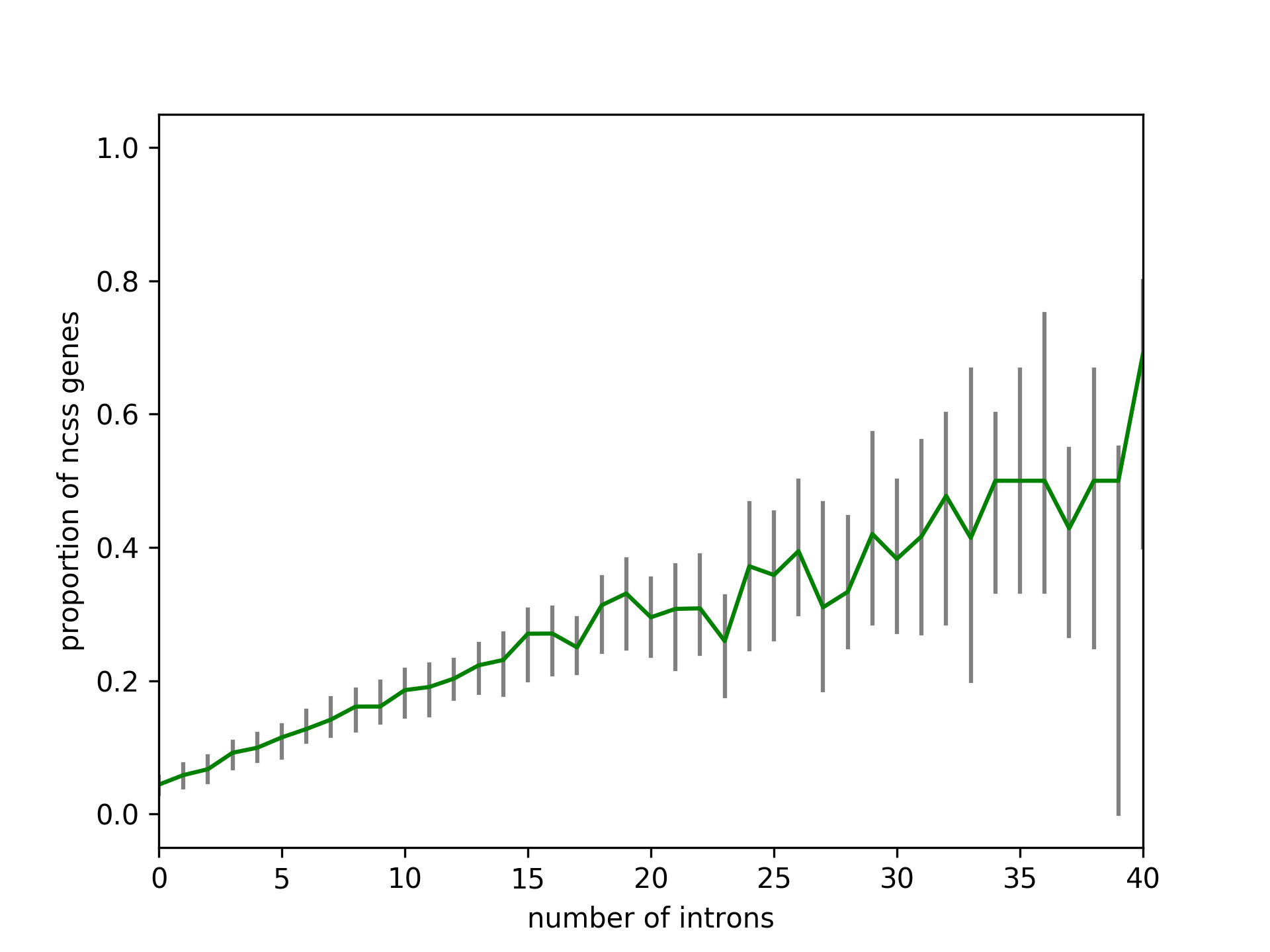

### Supplementary file 18

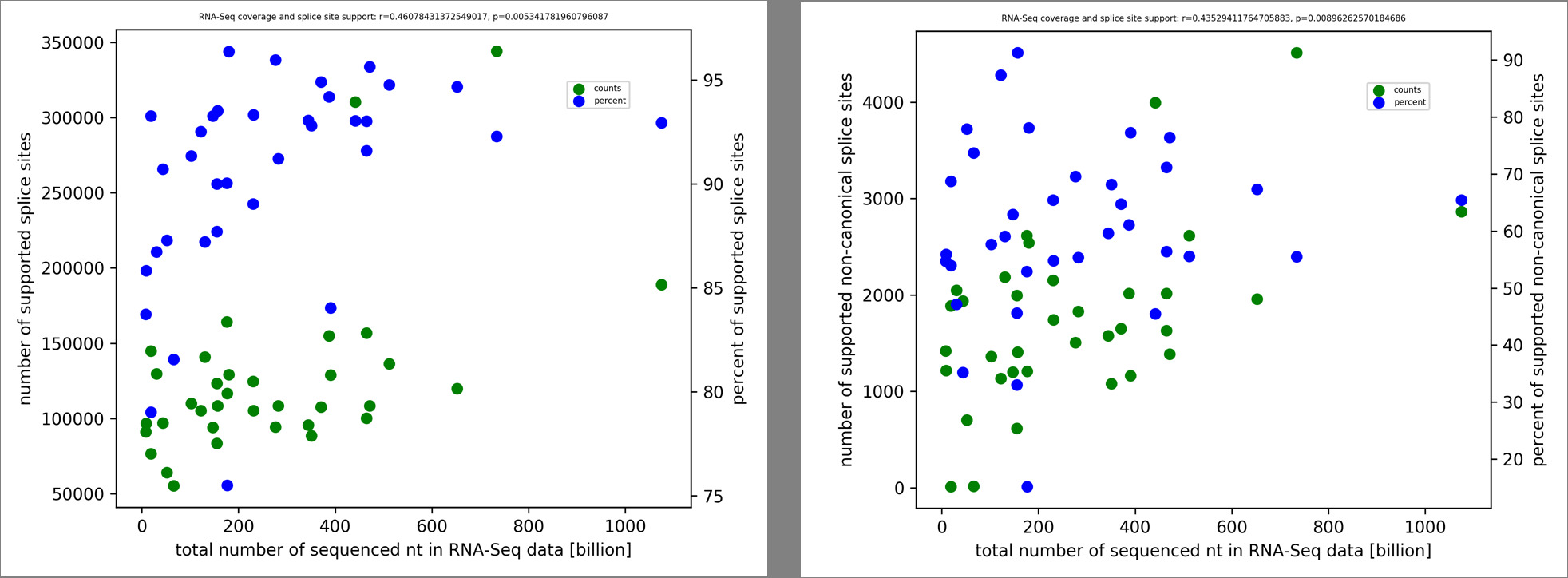
